# Human Mammary Cells in a Mature, Stratified Epithelial Layer Flatten and Stiffen Compared to Confluent and Single Cells

**DOI:** 10.1101/2020.03.08.982660

**Authors:** Hyunsu Lee, Keith Bonin, Martin Guthold

## Abstract

The epithelium forms a protective barrier against external biological, chemical and physical insults. So far, AFM-based, micro-mechanical measurements have only been performed on single cells and confluent cells, but not yet on cells in the physiologically relevant, mature epithelial layer.

Using a combination of atomic force, fluorescence and confocal microscopy, we determined the changes in stiffness, morphology and actin distribution of human mammary epithelial cells (HMECs) as they transition from single cells to confluency to a mature epithelial layer.

Single cells have a tall, round (planoconvex) morphology, have actin stress fibers at the base, have diffuse cortical actin, and have a stiffness of 1 kPa. Confluent cells become flatter, basal actin stress fibers start to disappear, and actin accumulates laterally where cells abut. Overall stiffness is still 1 kPa with two-fold higher stiffness in the abutting regions. Cells in an epithelial layer are flat on top and seven times stiffer (average, 7 kPa) than single and confluent cells. Epithelial layer cells show strong actin accumulation in the regions where cells adjoin and in the apical regions. Stiffness is significantly enhanced in the regions of adjoining cells, compared to the central regions of cells.

Physiologically, this previously unrecognized, drastic stiffness increase may be important to the protective function of the epithelium.

## Introduction

### Importance of biomechanical properties of cells

Over the last two decades, microscopic biomechanical analysis techniques, such as micropipette aspiration^1^, AFM indentation^2^, and magnetic and optical tweezers^3,4^, have progressed to the point where it is possible to study the mechanical properties of individual cells with relative ease. These studies have shown that cell mechanical properties are different for different cell types. The stiffnesses measured for isolated, cultured cells range approximately from 0.1 kPa to 40 kPa^5^, and the stiffness of cells correlates with biological function and the mechanical properties of tissue of origin. For example, neurons, constituents of the brain, which is one of the softest tissues, are soft with cell stiffness ranging from 0.1-2 kPa^6^. On the other hand, cardiac myocytes, which make up the cardiac muscle, are very stiff with elastic moduli in the 35-42 kPa range^7^. These studies also indicated that mechanical properties of cells correlate with their microenvironment and with cellular processes, including cell division^8^, adhesion^9^, migration^10^, motility^11^ and differentiation^12^. Moreover, several human diseases closely correlate with abnormal stiffening of cells, e.g., asthma^13^, vascular disorders^14^ and aging^7,15,16^; or softening of cells, e.g., cancer^17–20^. Therefore, investigating the mechanical properties of cells can provide valuable insights into various cellular processes, disease and cancer progression, and they may be used as a potential biomarker.

### Actin

Filamentous actin (F-actin) is a semi-flexible polymer, which is assembled from the monomeric, globular form of actin (G-actin). Actin filaments are highly dynamic and responsible for many cellular processes, including cytokinesis, changes in cell shape, cell motility, and intracellular trafficking. Additionally, actin filaments play an important role as a mechanotransducer, translating external physical forces into biochemical signals and leading to various cellular responses^21–24^. Actin filaments are organized into two types of main structures, stress fibers and cortical actin filaments, which play different mechanical roles within cells. Stress fibers, together with non-muscle myosin II, generate mechanical forces during movement, and they are responsible for the formation and maintenance of cell-to-cell or cell-to-extracellular matrix adhesions^25–27^. Cortical actin filaments form 50-200 nm thick networks underlying the inner surface of the plasma membrane.^28^ They endow the cell with its mechanical integrity and contractility, which is important for cell shape and deformability. The components of the cytoskeleton – actin filaments, microtubules and intermediate filaments, the plasma membrane, the nucleus, and other cellular organelles are possible determinants of the mechanical properties of cells. Numerous studies, however, have shown a close correlation between actin filaments and cell stiffness, indicating that actin filaments are the primary contributor to cell mechanical properties. This is likely due to their higher-order structures and interaction with crosslinkers and other actin binding proteins. Cells treated with reagents that inhibit actin polymerization, such as cytochalasin D and latrunculin B, showed significantly reduced cell stiffness^29–33^. Conversely, cells treated with nocodazole, which interferes with microtubule polymerization, showed insignificant changes in cell stiffness^29,32–34^. Cancer cells are softer and easier to deform than their healthy counterparts^20^. It has been proposed that the lower stiffness of cancer cells is attributed to a reduction in the amount of actin filaments or an increase in disorganized actin structures^35–37^.

*In vivo*, cells continuously exchange signals with their neighboring cells, and they are linked together by cell-to-cell junctions, which are responsible for regulating tissue homeostasis and integrity^38,39^. One of the major cell-to-cell junctions is the adherens junction where cadherin receptors serve as adhesion molecules, and actin filaments interact with them via the catenin protein complex, which form a bridge between the cytoplasmic domains of cadherins and actin filaments. The cadherin-actin interactions also play a role in the reorganization of actin filaments during development of adherens junctions. For example, actin filaments associated with e-cadherins are perpendicularly oriented to the plasma membrane at early stages of development of adherens junctions of epithelial cells. In contrast, actin filaments align parallel to the cell borders in mature epithelial sheets.^40,41^

### Motivation

As outlined above, the relevance of the mechanical properties of cells to a deeper understanding of key cellular processes was recognized over the last several years and a significant amount of work has been done determining the mechanical properties of different cell types. Initially, many studies concerning cell stiffness were performed on single, isolated cells, and the influence of confluency, cell packing, and cell-to-cell contact on cell stiffness has been underappreciated^35,36^. Recently it has been recognized that confluency and cell packing also need to be considered. Cell-cell interactions via adherens junctions underlie long-range correlations in cell stiffness^42^. Human mammary epithelial cells inside a colony (confluent layer) are stiffer than isolated cells^43^. The stiffness of MDCK II cells depends on confluency and cell size^44^. In most of these studies confluent cells were slightly stiffer, by a factor of two or less, than isolated cells^42–46^. However, very few studies, if any, were performed on mature epithelial layers that had been grown for several days, which is the physiologically most relevant structure of epithelial cells.

### Current work

Using a combination of atomic force microscopy, fluorescence microscopy and confocal microscopy, we investigated the stiffness, overall morphology and actin distribution of human mammary epithelial cells (HMECs) as they transition from single cells to confluency to a mature epithelial sheet conformation. We found that morphology, actin distribution and stiffness change significantly during this transition. Single cells have a tall, round morphology, with actin stress fibers at the base, diffuse cortical actin within the cell, and a stiffness of about 1 kPa. As cells become more confluent, they become flatter, actin stress fibers at the base start to disappear, and actin accumulates on the side where cells adjoin neighboring cells. The overall stiffness of these cells is similar to single (isolated) cells; however, with higher stiffness values in the adjoining regions where actin accumulates. Cells in an epithelial layer are flat at the top (apex), consistent with the physiologically typical, stratified, cuboidal epithelium form of HMECs. These cells are several times stiffer than single and confluent (but not yet packed) cells. Epithelial layer cells show strong actin accumulation in the region where cells adjoin and in the apical region. Stiffness is not uniform, as it is significantly enhanced in the regions of adjoining cells, compared to the central region of the cell. This previously unrecognized increase in cell stiffness could be physiologically important since epithelial cells form a key barrier to protect the body from biological, chemical, and mechanical insults.

## Materials & Methods

### Cell culture

Primary human mammary epithelial cells (HMECs; Lonza, Basel, Switzerland) were cultured in T-75 flasks in mammary epithelial basal medium (MEGM basal medium CC-3151; Lonza) that was supplemented with bovine pituitary extract, insulin, hydrocortisone and recombinant human epidermal growth factor (MEGM Bullet Kit CC-4136; Lonza). Cells were maintained in an incubator under standard conditions (36.5 °C, 5% CO_2_). HMECs were used within 6 passages.

50 mm glass bottom dishes (FluoroDish; World Precision Instruments, Sarasota, USA) were used for the imaging and indentation experiments. Collagen type IV (100 μg/ml in Dulbecco’s Phosphate-Buffered Saline; Sigma-Aldrich, St. Louis, USA) and laminin (100 μg/ml in DPBS; Sigma-Aldrich) were mixed at a 1:2 ratio to coat the dishes with 1 µg/cm^2^ of collagen IV and 2 µg/cm^2^ of laminin. The coating solution was placed into the dish, and the dish was incubated for 1 hour. The remaining solution in the dish was removed, then the dish was rinsed with sterile, deionized water. The dish was dried for 15 minutes under UV light before adding cells.

HMECs at 70-80% confluency were reseeded onto the prepared 50 mm dishes and cultured in the incubator for 1 to 3 days before the atomic force microscope and other imaging experiments were performed. Cell reseeding densities of 500 cells/cm^2^ (low), 10,000 cells/cm^2^ (medium) or 40,000 cells/cm^2^ (high) were used to control the confluency state of the cells, resulting in isolated cells, confluent cells and epithelial cell layers, respectively.

### Atomic Force Microscopy

An atomic force microscope (MFP-3D-BIO; Oxford Instruments, Abingdon, UK) combined with an inverted optical microscope (X73; Olympus, Tokyo, Japan) was used to carry out topographical imaging and nanoindentation of live cells. A cell culture dish was mounted on the petri-dish heater stage and the cells were maintained in culture medium at 36.5°C during the AFM experiments. Each session for the AFM measurements was conducted within 3 hours of removal from the incubator to avoid effects of pH changes of the culture medium on cells. A soft AFM probe (Biotool cell XXL, nominal spring constant k = 0.1 N/m, resonance frequency f = 50 kHz; Nanotools, Munich, Germany) was used for topographical imaging of live cells. The probe has a long conical carbon tip, which is critical to obtain a good image due to the height of the cell compared to the tip height; cantilevers with shorter tips may touch the cell with the side of the cantilever, rather than with the tip. Topographic images were obtained in AC mode with a 0.5 Hz scanning rate over an 80 × 80 µm^2^ area. A tipless, rectangular silicon nitride AFM cantilever (HYDRA6R-200NG-TL, nominal spring constant k = 0.035 N/m, resonance frequency f = 17 kHz; Applied NanoStructures, Mountain View, USA), to which a 5.3 µm diameter fluorescent melamine microsphere (Microsphere-Nanospheres, Cold Spring, USA) was attached, was used for the cell indentation experiments. The protocol to attach the microsphere is described next.

A clean glass coverslip was used as a substrate for the 5.3 μm microspheres and glue. 20 µl of microspheres in 2.5% aqueous suspension were deposited on the glass coverslip. A paper wipe was used to absorb and dry the water after the spheres sank to the coverslip surface, which occurred in a few minutes. Most spheres formed clusters, but some isolated spheres existed on the coverslip; these isolated spheres can be picked up by a cantilever. Using a pipette tip, a small amount (approximately 20 µl) of mixed, two-part marine epoxy glue was transferred onto the same coverslip. Using another coverslip, the glue was spread to a thin layer close to where the microspheres were deposited. The glue was dried for 20 minutes before moving on to the next step; drying for this period increased the viscosity of the low viscosity glue to a viscosity level appropriate for the following dip-pen nanolithography step. The AFM mounted on the optical microscope was used as a nanomanipulator for fine positioning, and the micrometers for the movement of the sample stage and the AFM head were used for coarse positioning. After putting the coverslip on the sample stage, the AFM cantilever was positioned in the middle of the field of view of the 40x lens. The laser was focused on the end of the cantilever, as is standard, and the deflection signal was monitored during the following processes to avoid breaking the cantilever. The edge of the spread glue on the coverslip was moved under the end of the cantilever. A small amount of the glue was applied to the end of the cantilever by lowering the AFM head manually while observing the cantilever through the optical microscope. After applying the glue, the cantilever was retracted to a safe distance above the surface (> 20 µm) and then an isolated microsphere was positioned under the cantilever. Using the AFM software, which can control the position of the piezo scanners with nm resolution, the cantilever was finely positioned above the microsphere. The cantilever was lowered by the z-piezo scanner until it contacted the microsphere. A few seconds after the contact, the cantilever was retracted and taken out of the AFM head. The cantilever with the microsphere was cured overnight under ambient conditions. A representative bead-modified cantilever is shown in Fig. 1.

**Figure 1.**
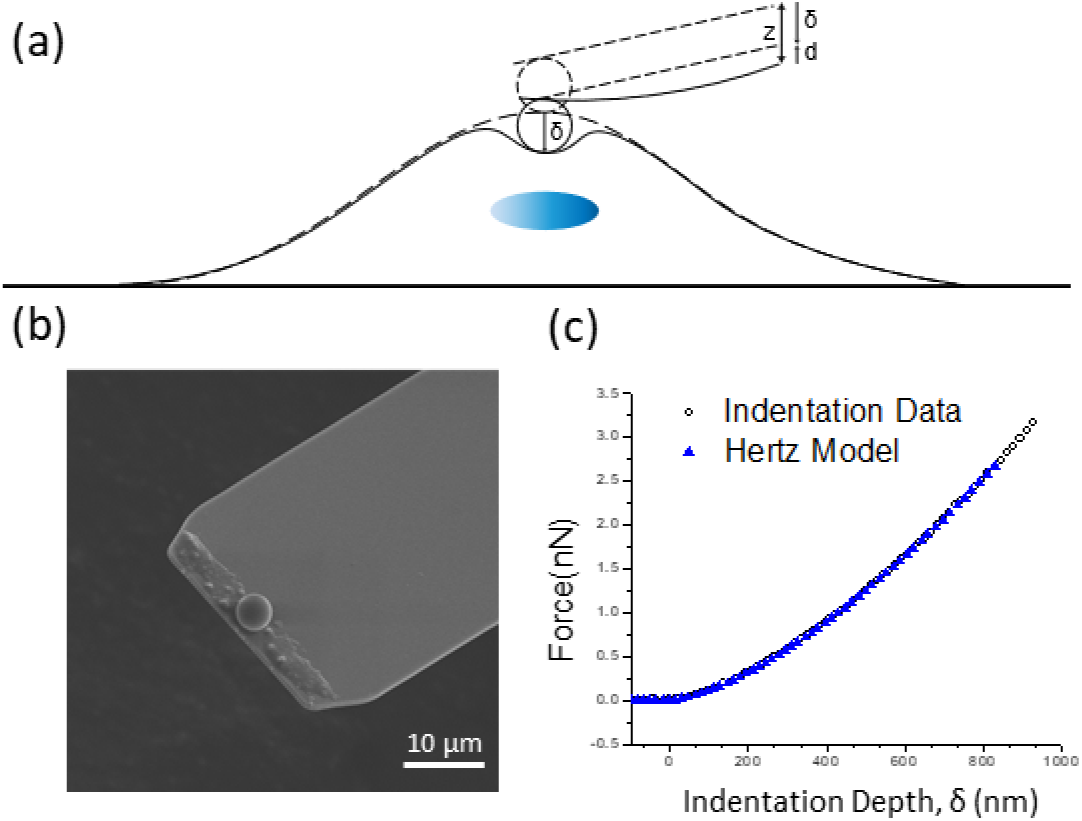
(a) Schematic of experimental indentation set-up. A cell is indented by an AFM cantilever with a spherical probe. The vertical position of the cantilever is moved by z; the cantilever bends by d; the cell is indented by δ; z = d + δ. (b) Top-view SEM micrograph of an AFM probe with an attached 5.3 μm microsphere. (c) Example of a force vs. indentation curve obtained by atomic force microscopy. The indentation data was fit using the Hertz model (equation 1).

To measure the spring constant of an AFM probe, we used the ‘GetReal’ automated probe calibration function in the Asylum Research software, which is based on the thermal noise method^47^ and Sader’s method^48^. The software first determines the spring constant, using Sader’s method with Q factor, the resonance frequency from the thermal noise spectrum, and the dimensions of a cantilever. Subsequently, GetReal determines a parameter, the so-called inverse optical lever sensitivity (InvOLS), which is needed to convert the photo diode signal in volts to displacement of the cantilever in nanometers, using the thermal noise method and the spring constant calculated in the previous step. However, we used the InvOLS determined by a different, more direct way. Specifically, by measuring the slope of the deflection voltage vs. z distance curve on a hard surface. There was usually a 10-20 % discrepancy between this slope method as compared to the thermal noise method. Since the latter slope method is more direct, we used that value.

The following equation, derived from the Hertz model, was used to determine the Young’s modulus of each cell.

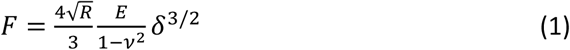

Here *F* is the indentation force, *R* is the bead radius, *E* is the Young’s modulus, ν is the Poisson’s ratio of a cell (taken to be 0.5), and *δ* is the indentation depth. Three to five indentations were carried out on the center of each cell to obtain force versus indentation depth curves. Since the Young’s modulus of cells depends on the loading rate and indentation depth, the speed of the indentation was fixed at 5 µm/sec and the indentation depth was maintained in the range of 0.6 to 1 µm. The obtained force curves were processed using the built-in Hertz model fitting tool of the Asylum AFM software. The Young’s modulus values of the curves were arithmetically averaged for each cell. A representative indentation curve and fit, according to the Hertz model, is shown in Fig. 1(c).

A stiffness map, also called a force-volume map, consisting of 32 × 32 pixels within an 80 × 80 µm^2^ area, was acquired by taking individual force versus indentation curves on each pixel. Like the single indentation, the indentation depth and loading speed for a stiffness map on each pixel were set in the range of 0.6 to 1 µm and 5 µm/sec, respectively. It took approximately 18 minutes, which is short enough to avoid drifts caused by cell movements, to obtain a stiffness map with these conditions. The Young’s modulus of each pixel was determined by fitting the Hertz model for the 5.3 µm microsphere to the individual curves, using the analysis tools of the AFM software.

### Fluorescence microscopy

For fluorescence imaging of F-actin and nuclear structures of live cells, F-actin filaments and the nuclei were labeled with 1 µM of SiR-actin (Cytoskeleton, Denver, USA) and 1 µg/ml of Hoechst 33342 (Invitrogen, Carlsbad, USA), respectively. Each of the dyes, dissolved in DMSO, were directly added to the culture medium, and the cells were incubated for 30 minutes prior to fluorescence imaging.

Epifluorescence images correlating to the AFM topographic images were taken by the X73 Olympus inverted microscope, which is situated under the AFM, with a 60x oil immersion lens (numerical aperture, NA = 1.3) and a Hamamatsu digital camera (C11440, ORCA-Flash2.8, Hamamastsu, Hamamatsu, Japan). Filter sets for DAPI and Cy5 channels were used for Hoechst 33342 and SiR-actin, respectively.

For confocal fluorescence imaging, a confocal microscope (LSM 880; Carl Zeiss, Oberkochen, Germany) in Airy-scan mode with a 63x oil immersion lens (NA = 1.40) was used. A 405 nm diode laser and a 633 nm He-Ne laser were used as excitation lasers for Hoechst 33342 and SiR-actin. Each image was acquired by scanning twice with the two lasers in turn. Cross-sectional views consisting of 40-60 layers were acquired by single-line scanning which can be done in a few minutes.

## Results

One of the key functions of epithelial cells is to form a protective barrier between the inside of the body and the outside world. The overall goal of our work was to characterize the global morphology, stiffness and actin distribution of human mammary epithelial cells as they progress from single cells, through confluency, to a mature, epithelial layer.

### Global morphology and actin formation of HMECs at different degrees of confluency

AFM and fluorescence images were taken to determine how cellular morphology and F-actin structure change with increasing cell confluence (Fig. 2). AFM deflection images (error signal) indicate the topography of single HMECs, confluent HMECs, and HMECs of epithelial layers (Fig. 2 a-c). Cell areas tend to decrease as cells reach confluence and then form the basal layer of the epithelium. However, additional cells that form the apical level (second layer) of the epithelium had the largest area compared to single, confluent cells and basal epithelial layer cells. The cross-sectional profiles (Fig. 2 b), corresponding to the white lines in Fig. 2a, show the height differences between the lowest and the highest topographical features. The measured height differences for the single cell, confluent cells and the epithelial layer shown in Fig.2 are 8.5 μm, 2.5 μm and 300 nm, respectively. This indicates that cells become flatter as they reach confluence and form the epithelium. We took epi-fluorescence microscope images of actin filaments stained with SiR-actin of the cells corresponding to the AFM images (Fig 2. g-l). Using a 60x oil immersion lens (NA = 1.3) and shallow depth of field, it was possible to distinguish actin filaments in the apical focal plane (Fig. 2 g-i) and basal focal plane (Fig. 2 j-l). In the apical plane of the single cell (Fig. 2g), the actin distribution is diffuse, with no apparent, distinct actin filaments. The image of the apical focal plane of the confluent cells (Fig. 2h) shows thin cortical mesh actin filaments in the middle of cells and a strong, blurred distribution of circumferential actin. Some of the circumferential actin is in focus and some out of focus, indicating that it is vertically distributed along the entire height of the cell. In the apical focal plane of the epithelial layer cells (Fig. 2I), we observed dense, distinct, circumferential actin filaments and filaments that stretched out to span cell-to-cell contact regions. Note that the apical plane has a smaller number of cells than the basal plane. Thick and dense dorsal stress fibers were observed in the basal focal plane of the single cell (Fig. 2j), which indicates that the cell was actively migrating toward the upper left of the image. The image in the basal focal plane of the confluent cells (Fig. 2k) shows that thinner, sparse, and unidirectionally aligned stress fibers and actin filaments surrounding the cells were formed, indicating a more stationary and collectively migrating cell. Overall, in the basal focal plane of the cells in the epithelial layers (Fig. 2l), the amount of stress fibers is markedly decreased, and the stress fibers are randomly oriented, indicating non-migrating, spatially stable cells.

**Figure 2.**
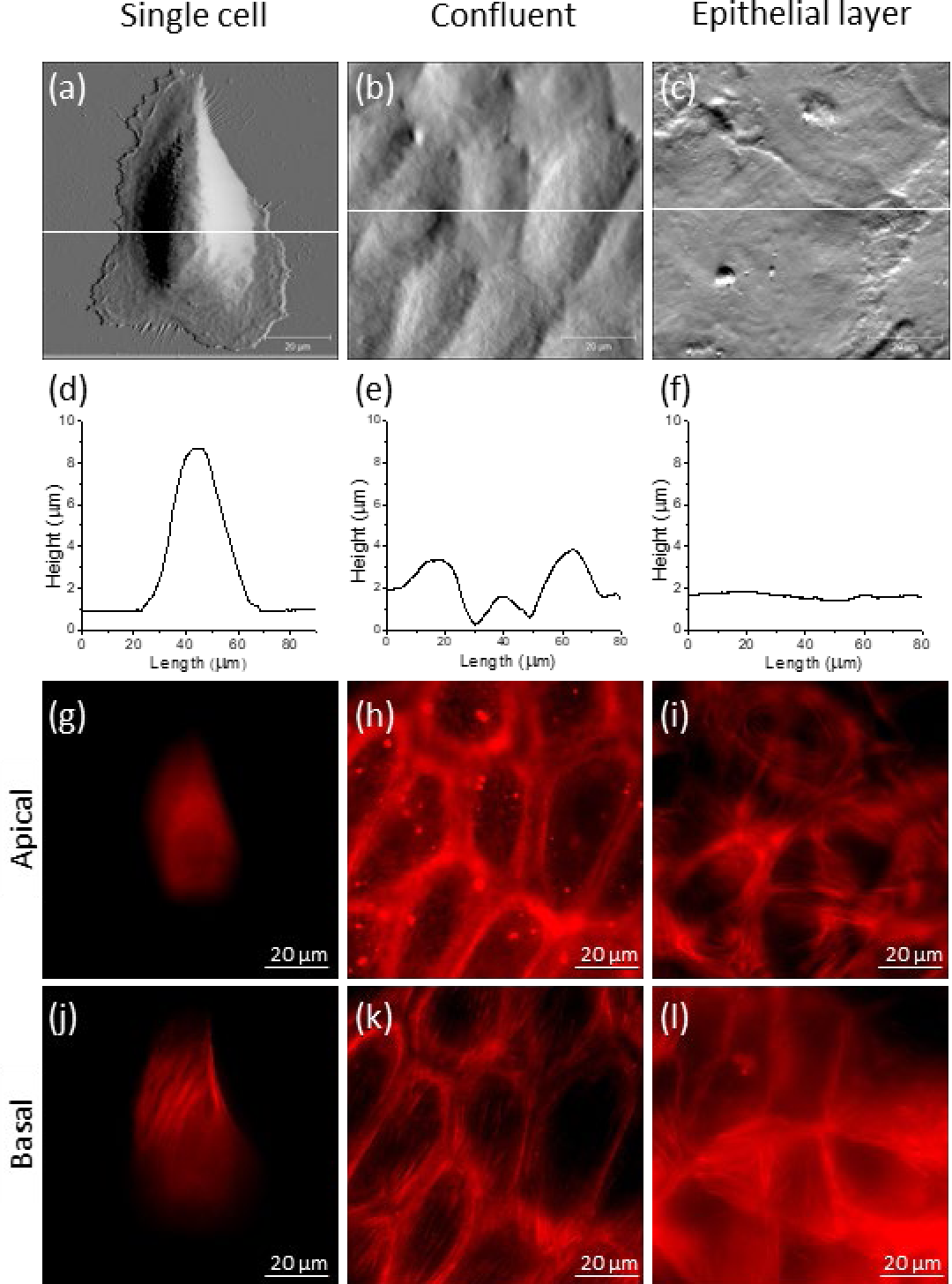
AFM deflection images of HMECs in (a) single, (b) confluent, and (c) mature epithelial layer states. (d-f) Cross-sectional line profiles of the AFM deflection images corresponding to each of the white lines in (a-c). Fluorescence images of actin filaments stained with SiR-actin in the (g),(h),(i) apical and (j),(k),(l) basal levels of the cells.

### Young’s modulus of individual cells measured with an AFM

The first two histograms (Figs. 3a & b) show the distributions of Young’s moduli measured in the central region of individual cells in the single and confluent states, and the third histogram (Fig. 3c) is the distribution of Young’s moduli measured on the top layer of the epithelium. Note that the indentation locations on the top layer were randomly chosen due to the ambiguity in determining the borderlines between cells that is a result of the continuous and flat morphology as shown in Fig. 2c. Also, note that the x-axis scales of Figs. 3a & b are the same, while the scale of Fig. 3c is much larger. The width of the distribution of single cell moduli (Fig. 3a) is slightly wider than that of confluent cells (Fig. 3b), while the averaged Young’s modulus, E, of single cells and confluent cells are equal within error, with E = 1.01 ± 0.52 kPa (N = 183) and 1.09 ± 0.51 kPa (N = 199), respectively. According to a Student’s t-test analysis, the two distributions are not statistically different (p > 0.5). The width of the distribution of Young’s moduli of the epithelium (Fig. 3c) is much wider than the other two distributions, ranging from 0.37 kPa to 37 kPa, and the averaged Young’s modulus of the epithelium increased about seven-fold to 7.09 ± 5.8 kPa (N = 717). Compared to single and confluent cells, the increase in Young’s modulus is statistically significant (p < 0.001).

**Figure 3.**
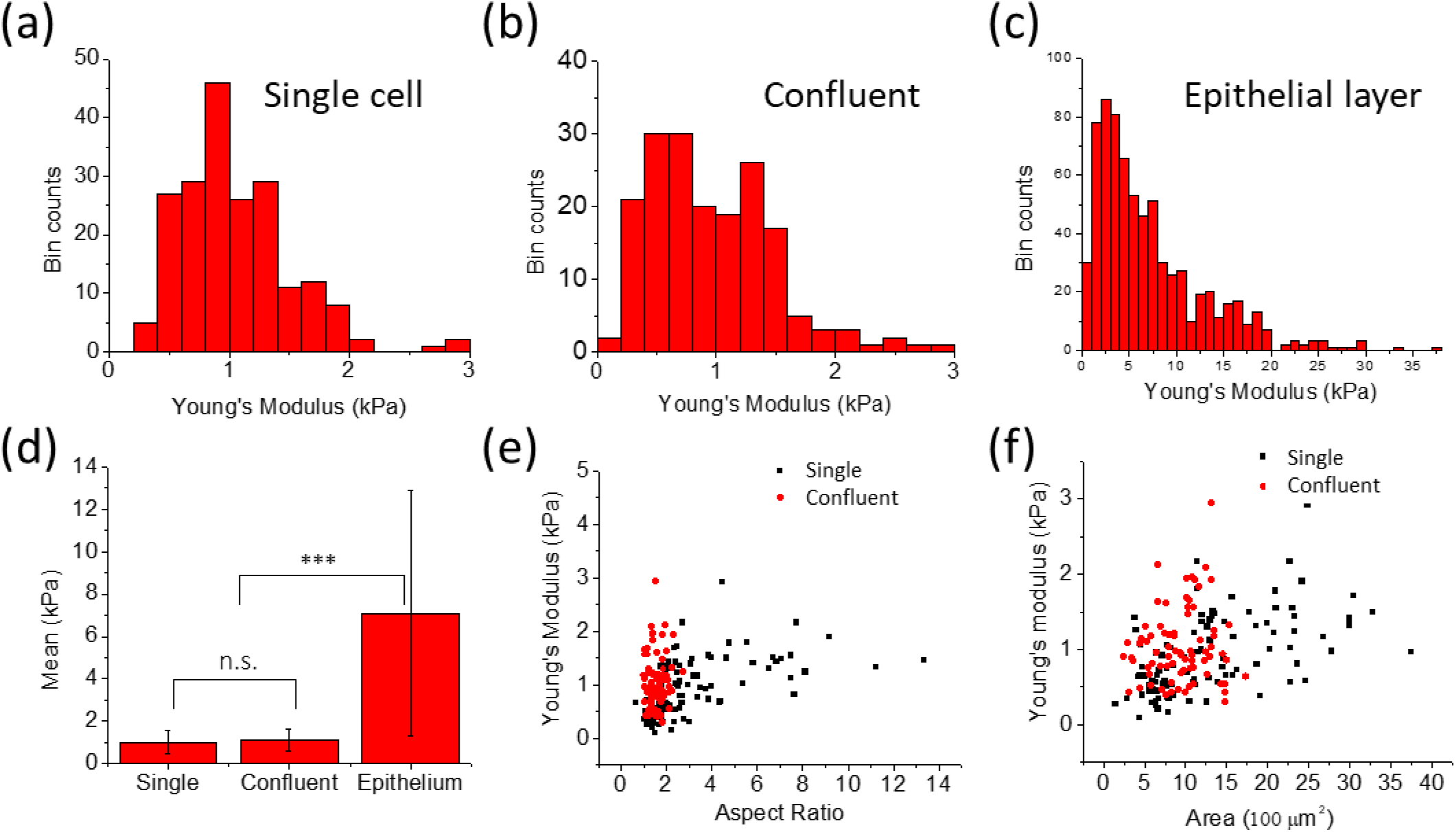
Histograms of the Young’s moduli obtained from single AFM indentations of single cells (a), confluent cells (b), and cells in an epithelial layer (c). (d) Mean Young’s modulus plots from the histograms. (e) Young’s modulus vs aspect ratio plot. (f) Young’s modulus vs area. Pearson’s correlation coefficient analysis results in correlations for single cells between the Young’s modulus and cell aspect ratio (R = 0.51) and area (R = 0.47). The same analysis on confluent cells showed they were uncorrelated for both aspect ratio (R = 0.06) as well as for area (R = −0. 06).

To determine if overall cell area and shape affect stiffness, we plotted the Young’s modulus vs. aspect ratio and Young’s modulus vs. cell area (Fig. 3e & f). Note that the data used for these plots are sub-sets of the histograms, but the data for the epithelium was not included in these plots because the modulus magnitudes differ significantly (see discussion). The aspect ratios of the single cells ranged from approximately 2 to 14, and the Young’s modulus of single cells tends to increase with higher aspect ratios. On the other hand, the distribution of the aspect ratios of the confluent cells was localized around 2 with the Young’s modulus ranged between 0.3 and 2 kPa. Single cell areas ranged from 250 µm^2^ to 3800 µm^2^, and the Young’s modulus tends to increase as cell area increases. The areas of the confluent cells did not exceed 1800 µm^2^, and there was no correlation between cell area and stiffness. Taken together, although the Young’s moduli of both single and confluent cells ranged between 0.3 kPa and 3 kPa and the two averaged values are very similar, single cells saw their stiffness (force/area) rise as each cell stretched out over a larger region. When comparing cells smaller than 1700 µm^2^ (see Fig. 5f), confluent cells were slightly stiffer than single cells in this small cell area regime.

**Figure 4.**
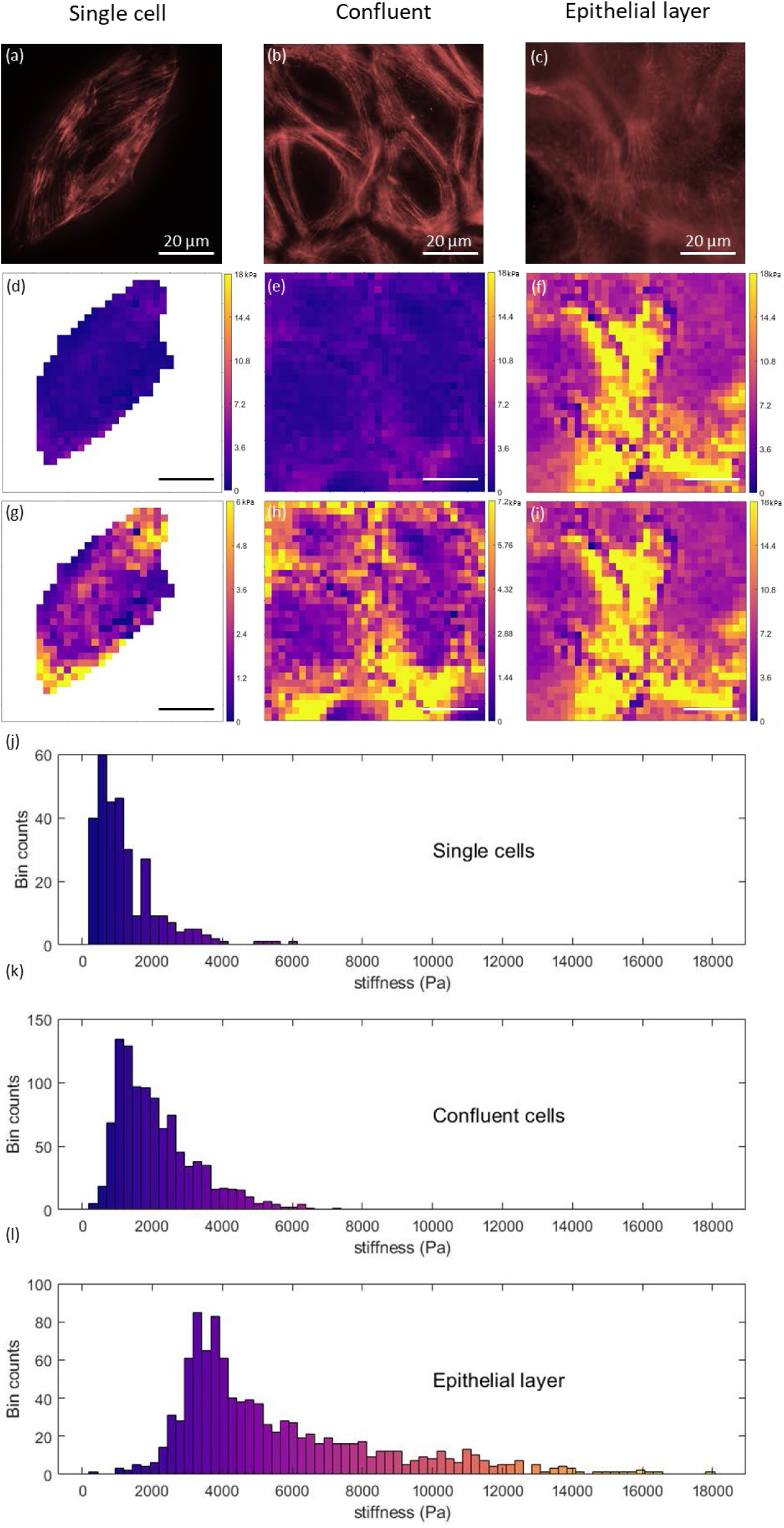
(a),(b),(c) Fluorescence images of actin filaments of HMECs in single, confluent and mature epithelial layer states. (d),(e),(f) Stiffness maps using the same Young’s modulus scale. (g),(h),(i) Stiffness maps using a normalized scale (not the same across images). (j), (k), (l) Histogram plots for each stiffness map. The histogram color map has the same values as the color map for images in (d), (e), (f).

**Figure 5.**
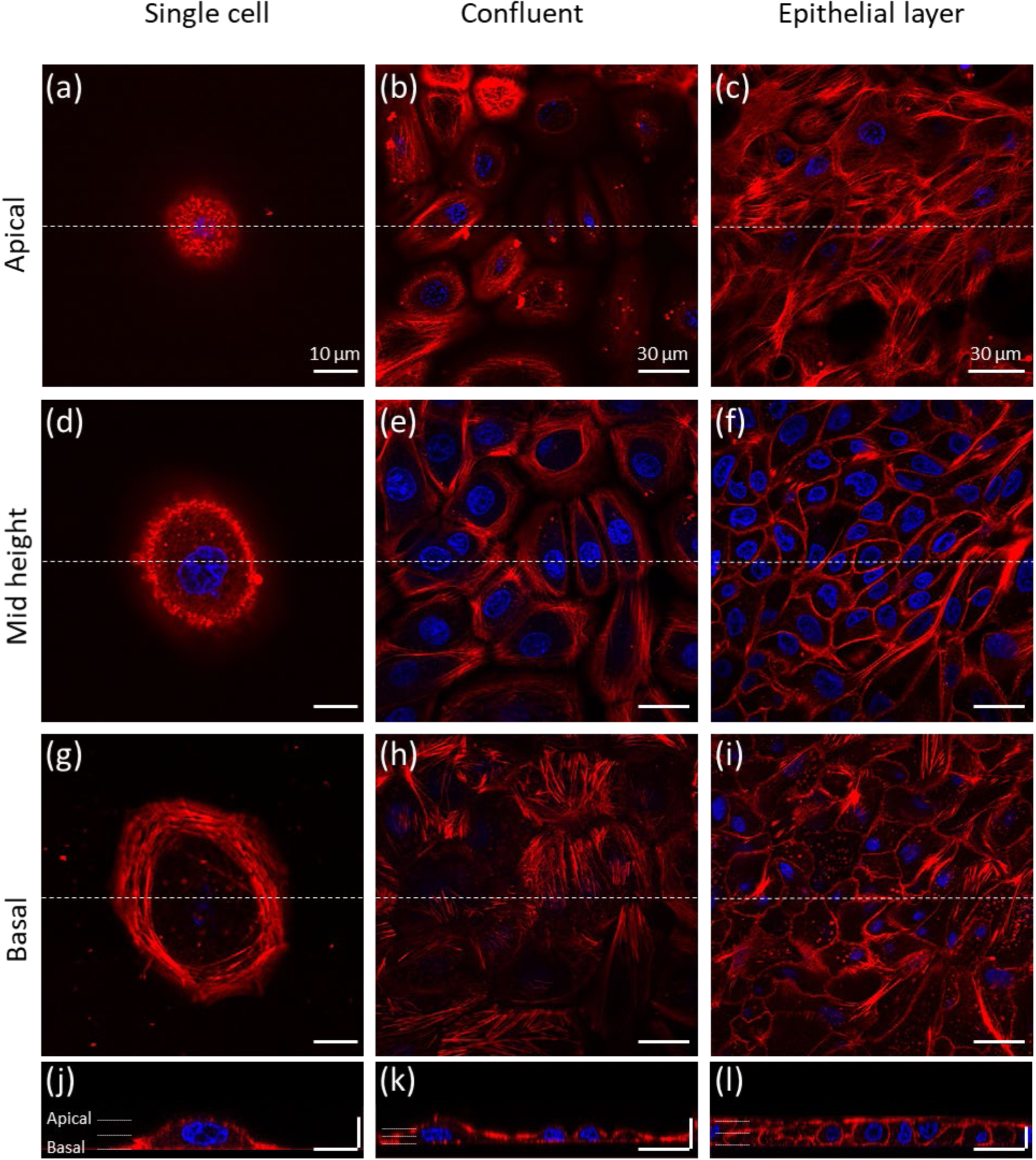
Confocal microscope images of F-actin (red) and nuclei (blue) of HMECs taken in (a), (b), (c) apical, (d), (e), (f)mid-height, and (g), (h), (i) basal focal planes. (j), (k), (l) Cross-sectional view. Horizontal scale bars are 10 μm for single cells and 30 μm for confluent and epithelial layer cells; each vertical scale bar is 10 µm.

### Stiffness Maps of Cells

To investigate the correlation between actin filament formation and cell stiffness, fluorescence microscope images of the stained actin filaments of cells were taken, followed by force mapping of the same cells with the AFM (Fig. 4). It can be seen that single cells have the lowest stiffness, and there were no distinct features in the stiffness map that correspond to actin formation. This is likely because there are no actin filaments directly contributing to the stiffness except for cortical actin; actin is found in the basal stress fibers and the diffuse cortical actin. The reason why the edges of the single cell were stiffer is due to the thin film effect – the stiffness of the substrate is ‘felt’ when measurements are done on thin films. In the case of confluent cells, the Young’s modulus of the central region of the cells with no distinct actin formation was similar to that of a single cell. However, the Young’s moduli of the regions with circumferential actin filaments are twice as stiff as the central regions of the cell. In the case of the epithelium, cell stiffness markedly increased over the whole area. In the cell-to-cell contact regions of the epithelium, dense actin filaments are formed, and the stiffness was highest in these regions with values of 10-14 kPa. The images indicate that overall, the Young’s modulus tends to have higher values in regions where actin filaments are dense.

### F-actin Distribution via Confocal Microscopy

To investigate how actin filament formation changes due to increased cell-cell interactions as cells reach confluence, images of SiR-actin stained actin filaments were taken by confocal microscopy. Fig. 5 shows actin filament formation in the apical (Fig. 5a-c), mid-height (Fig. 5d-f) and basal planes (Fig.5g-i) of single and confluent cells, and the epithelial layer. In single cells, only the cortical actin mesh network, which underlies the inner surface of the plasma membrane, is observed in both the apical (Fig. 5 a) and mid-height (Fig. 5d) focal planes. In the basal plane of the single cell (Fig. 5g), extensive circumferential stress fibers were observed. Unlike Fig. 2j, in which dorsal stress fibers were seen in a migrating cell, there are no dorsal stress fibers seen here, indicating that this single cell was not migrating in a specific direction. The cross-sectional image of the single cell (Fig. 5j) shows a typical morphology of an adherent cell which has a planoconvex shape in the middle. In confluent cells, except for the central region of the cells, dense actin filament bundles surrounding the cells were observed in the apical (Fig. 5b) and mid-height (Fig. 5e) focal planes. In the basal focal plane of the confluent cells (Fig. 5h), randomly oriented stress fibers were observed, and stress fibers have mostly disappeared in some of the cells. In the cross-sectional images of confluent cells (Fig. 5k), cells remained roundish, but the actin filaments were prominent, underlying the inner surface of the plasma membranes close to the cell-cell contact regions.

Cells in the apical level of the epithelial layer (Fig. 5c) were observed to have actin formation intricately intertwined between the cells. In the mid-height plane (Fig. 5f), the actin filaments forming the boundaries between cells were observed. In the basal plane (Fig. 5i), most of the stress fibers disappeared and remained in the form of dots with actin at the cell boundaries. In the cross-sectional image of the epithelium (Fig. 5 l), it was observed that the cells were arranged in an array of cubes under the very flat, apical actin layer (see also Fig. S1 in the supplementary information, which shows additional projections through the cells from apical to the basal region illustrating the mature epithelial layer structure).

## Discussion

### Significance and summary of results

Physiologically, epithelial cells line surfaces and ducts in the body that may come into contact with the external environment. Their tasks include providing a protective biological, chemical and physical barrier; being able to respond to physical forces and disturbances; and being able to migrate and rearrange in response to external stimuli and insults. The mechanical properties of epithelial cells are, therefore, critical to their function. Moreover, they are also of interest to cancer researchers as many cancers start in the epithelium.

Human mammary epithelial cells (HMEC) form a stratified cuboidal epithelium, which differs from a simple epithelium in that it is multilayered. It is typically found in gland linings that are specialized in selective absorption and secretion, that need to withstand mechanical or chemical insults, and where cells might be abraded. Cell strata in these epithelia become flatter as the strata become more apical^49^.

Numerous studies have examined the mechanical properties of both single and confluent epithelial cells. However, few, if any, studies have examined the mechanical properties of cells in a mature epithelial layer, and how mechanical properties change as cells transition from single cells to an epithelial layer.

The goal of our work was to determine the changes in gross morphology, cell mechanical properties (stiffness) and actin distribution in human mammary epithelial cells as they transition from single cells to confluent cells to an epithelial layer configuration. We used the following methods to characterize the cells: AFM imaging, fluorescence imaging, AFM-based nanoindentation, AFM force mapping with a 5.3 μm micro-bead probe, and confocal microscopy. Our key findings were:

1. **Morphology**. Single, isolated cells are roundish (planoconvex) and tall (8.5 μm); confluent cells have a flatter, less tall morphology; and mature epithelial layers have a flat top surface (apical side).
2. **Average stiffness vs. cell morphology.** Single, isolated cells are comparatively soft (1 kPa); confluent cells, even though they are flatter, still have an average modulus of 1 kPa; mature epithelial layers are, on average, much stiffer (7 kPa). Here, stiffnesses were measured over the cell center.
3. **Stiffness vs. area.** The stiffness of single, isolated cells positively correlates with their aspect ratio and their area.
4. **Stiffness maps.** Force volume measurements (stiffness maps) with a resolution of 2.5 μm reveal the fine spatial structure of cell stiffness. For single cells, stiffness is mostly uniform across the cell. For confluent cells, the stiffness is higher by a factor of two in the region that adjoins other cells as compared to the central region of the cell. For packed cells in an epithelial layer of cells, the region adjoining other cells is also stiffer than the central region, and overall, these cells are significantly stiffer (factor of 7) than single or confluent cells.
5. **Actin distribution**. The actin distribution changes significantly during the single cell to epithelial layer transition. In single cells, actin is mostly seen as stress fibers in the basal plane (indicative of migrating cells) and as diffuse cortical actin on the apical side. In confluent cells, there are still some, but less pronounced stress fibers in the basal plane. Actin is starting to be seen in the region where cells adjoin. In cells that have formed an epithelial layer, basal stress fibers have largely disappeared. Actin is pronounced and distinct in a strong band in the regions where cells adjoin. Moreover, there are clear actin fibers and actin accumulation in the apical plane. There is an additional layer of cells, on top of the basal plane cells. These top cells are very flat, spread-out and actin-rich. A visual summary of these morphological observations is given in Fig. 6.

### Cell stiffness in context of current literature

Our results on the AFM-measured modulus of *single, isolated* cells largely agree with values reported in the literature; typical stiffness values are on the order of 1 kPa for epithelial cells for indentation speeds on the order of several μm/s^43–45,50^. Compared to other cell types, single epithelial cells are soft; they are slightly stiffer than the very soft neurons^6^ (0.4 kPa), but significantly softer than cardiomyocytes^7^ (35 kPa) and osteoblasts^51^ (8.3 kPa). The tall, roundish morphology we observed also agrees with the shape of single epithelial cells reported in the literature.^35,36^ Pronounced actin stress fibers at the basal plane indicate migrating cells, while the diffuse cortical actin distribution is indicative of a soft apical region.

**Figure 6.**
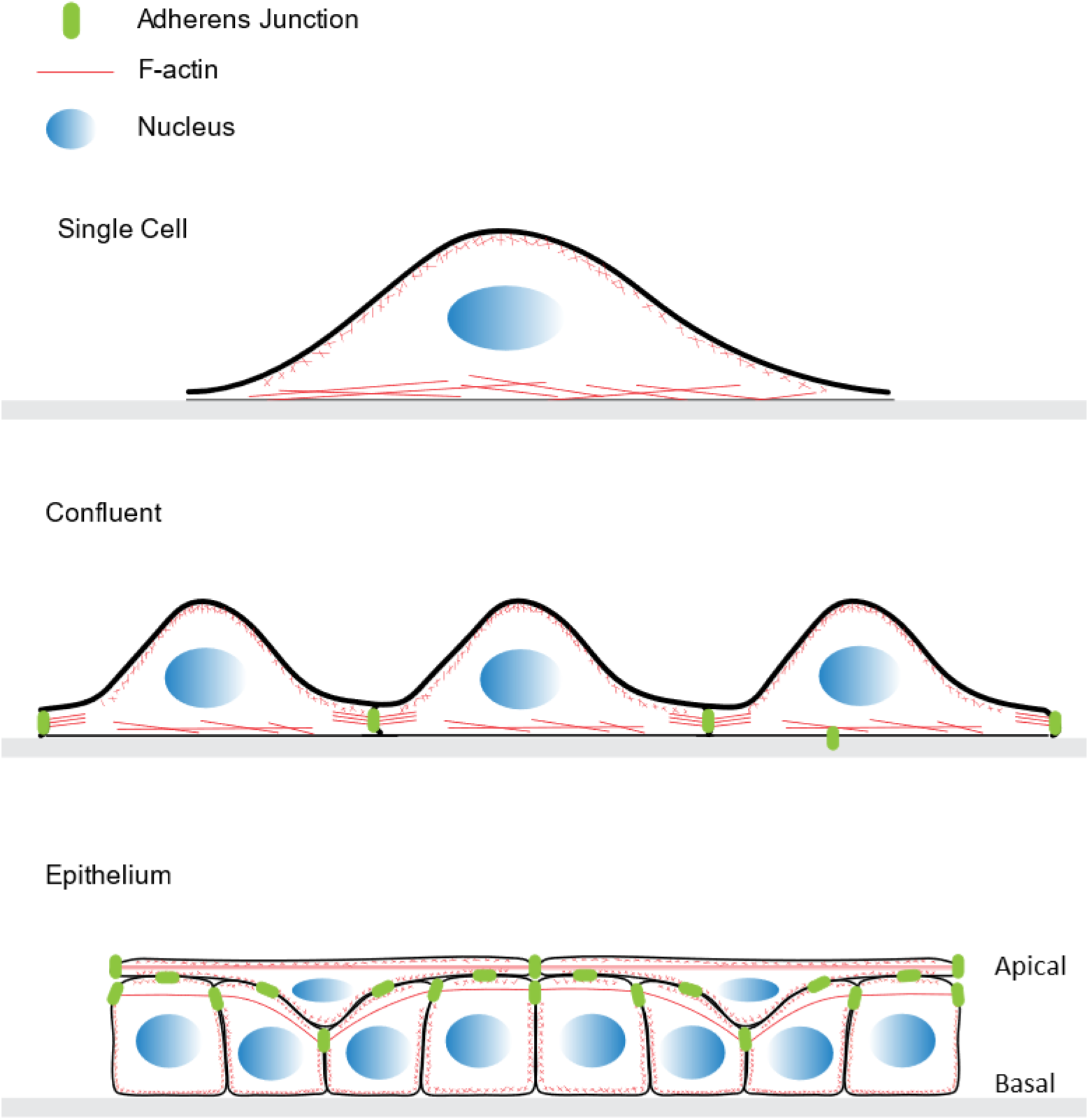
Schematic indicating progression of human mammary epithelial cell states. Isolated cells are roundish (planoconvex), tall (8.5 µm) and soft (1 kPa) with actin stress fibers at the basal plane. Confluent cells begin to flatten out; they are still soft (average, 1 kPa) with increased stiffness (2 kPa) and actin accumulation in the region of abutting cells, actin stress fibers at the basal plane start to disappear. Basal cells in an epithelial layer become cuboidal in shape, apical cells are very flat and actin-rich. Significant actin accumulates in the region of abutting cells. The average stiffness of the layer as measured from the top is significantly enhanced (7 kPa), with local stiffness in abutting regions reaching stiffnesses up to 30 kPa.

As cells become confluent, but not yet tightly packed and layered, cells are sensing each other, and they are forming adherens junctions^40,52^. In epithelial cells, the formation of adherens junctions triggers a decrease in RhoA activity and an increase in Rac1 and Cdc42 activity; it gives rise to slowed migration, a decrease in stress fibers, the formation of focal adhesions, and actin accumulation in the regions where cells abut^53,54^. Our observations agree with these changes, as shown in Fig. 5e & h. Concurrently, the stiffness in the peripheral cellular regions increases, while the stiffness in the center of the cell remains the same. The modulus we observed for confluent cells largely agrees with values reported in the literature for confluent epithelial cells^43–45^. As epithelial cells become confluent, it was observed that the modulus stays the same or slightly increases (by up to a factor of 2). The slight discrepancy between either staying the same or slightly increasing may be explained by the following factors. Different epithelial cell types or measuring conditions were used across the different experiments. Additionally, as can be seen in Fig. 4 h, the modulus varies depending on where on the cell it is measured; it is twice as stiff in the peripheral region. Finally, as our data show, a confluent layer is not the final stage of cell progression as cells further develop into an epithelial layer, which is several-fold stiffer. The somewhat higher stiffness values reported in the literature may have been on confluent cells that were transitioning to the epithelial layer stage.

Specifically, the following reports constitute some of the relevant key studies on the stiffness of epithelial cells. Guo et al.^43^ found that single, isolated HMECs (same cells we used) had a modulus of 1 kPa and they were 1.5 times softer than (sub-)confluent cells, while immortalized, tumorigenic, and metastatic cells (MDA-MB-231) did not (or only marginally) change in response to being surrounded by other cells. Similarly, Schierbaum et al.^55^ found that single mammary epithelial cells (MCF10A) are softer by a factor of 2 compared to cells in a confluent layer. Again, this change in stiffness was not seen in MCF7 and MDA-MB-231 cells. This lack of change in stiffness of the cancer cell lines with respect to confluency might relate to the lack of e-cadherin expression and a diminished F-actin belt. This, in turn, could result in deregulated cell proliferation as mechanical tension from adherens junctions and F-actin plays an important role in regulating cell proliferation.^22,56^ Absolute stiffness values were about 10 times larger in Schierbaum et al. as compared to Guo et al. and compared to the current study. This may be due to Schierbaum using sharp probes and somewhat different substrate and media conditions. Using dynamic, frequency-dependent, AFM-based mechanical measurements, Rother et al.^50^ found that malignant cells (tumorigenic MCF-7 and metastatic MDA-MB231) are typically softer than their benign counterparts (MCF-10A), and that malignant cells have a lower loss tangent (are more fluid-like). The Young’s modulus, as calculated from E = 2(1+ν)G’ is 4.5 kPa for MCF-10A cells (immortalized, benign cells similar to our HMECs) for indentation speeds and indentation depth similar to those in our measurements. In a series of publications, the Janshoff lab investigated the viscoelastic properties of Madin-Darby canine kidney cells, strain II (MDCK II) using two dynamic AFM-based indentation methods, oscillatory microrheology (OMR, tip performs small, 40 nm oscillations during indentation) and force cycle experiments (FCE, cyclic indentation curves).^44,57–59^ Their modulus values for confluent cells agrees with our values (∼1 kPa).

Notably, the mechanical properties of cells in stratified epithelial layers, which is the natural physiological state of mammary epithelial cells, has not been investigated in the past. We found that cells undergo a significant transformation as they arrange in a mature, epithelial layer. Cells become, on average, stiffer by a factor of 7. Stiffness is not uniform, as it is significantly higher in the regions where cells abut. Moreover, actin is seen in a strong, distinct belt in this region (especially visible in Fig. 5 f). There are, furthermore, flat, actin-rich cells on top of the lower cells, which is expected for HMECs as they form a stratified epithelial layer. This actin distribution and stiffening strongly suggest that thick actin fibers are a major determinant of cell stiffness (see *Actin* section below). It also suggests that this is physiologically important as the epithelial layer forms a barrier against external insults.

### Justification of Hertz model and error sources

We used the Hertz model of a hard sphere indenting an elastic plane to determine the Young’s modulus of cells (eq. 1). Although some approximations are made when this model is applied to cell indentations, the following considerations support its use: 1) It is straight-forward; 2) it has a relatively small, estimable error as compared to other, more complex methods (see below); 3) it allows easy comparison of cell stiffness data across labs; 4) the obtained results are physiologically meaningful; and 5) the obtained modulus can be easily compared with the moduli of numerous other materials. The error due to measurement geometry is small as the radius of the indenting sphere can be accurately determined and the cell surface is reasonably planar. The fits of this model to our experimental data are excellent. Another error comes from the fact that cells are viscoelastic objects, whereas the Hertz model treats cells as elastic objects. This error can be estimated from dynamic, AFM-based indentation methods that allow a separation of the viscous and elastic component of the modulus. Brückner et al.^60^ found that the elastic component dominates at indentations speeds of 2 μm/s, (similar to our 5 μm/s), since the power law coefficient β for epithelial cells is in the range of 0.25 to 0.3 (β = 0 means completely elastic deformation, β = 1 means Newtonian fluid with completely viscous deformation). Similarly, Schierbaum et al.^55^ determined a power-law exponent, β, of 0.1 and 0.12 for confluent and single MCF-10A cells, and 0.2 and 0.15 for MCF7 and MDA-MB-213 cells. These relatively low values of β justify using the elastic Hertz model to a first approximation; adding the viscous component into the elastic component (as is done when applying the Hertz model) results in an about 10-30% overestimation of the elastic component. Brueckner et al. found an elastic modulus (separate from the viscous component) of 0.6 kPa to 0.8 kPa for confluent MDCK II epithelial cells (indentation depth up to 1 μm). This is in very good agreement with our measurement of 1 kPa for the total modulus (elastic and viscous part), with the caveat that it was two different types of epithelial cells (MDCK II vs. HMEC).

There may be additional error sources; however, all of them are smaller than the factor of 7 increase in stiffness we observe for epithelial layer cells. Our indentations were small enough (600 nm to 1000 nm) to not ‘feel’ the nucleus or the hard glass substrate underneath the cell. In some instances when taking measurements over the thin, peripheral section of single cells, the shape of the indentation curve actually did deviate significantly from the Hertz model, resulting in very high stiffness values. In these instances, the substrate stiffness was measured, and these data were eliminated. Typically, the probe-sample contact point in the force-indentation curves was well defined (Fig. 1b), indicating that these cells have a thin glycocalyx. Our experiments were all carried out on a hard, functionalized substrate (collagen- and laminin-treated glass) in temperature-controlled (36.5°C) and pH-controlled media. Recently, it was shown that cell stiffness is somewhat influenced by the stiffness of the substrate, as cells grown on stiff substrates cells are about 1.5 to 3 times stiffer than cells on soft substrates.^61,62^ However, the stiffening factor due to the transition from single/confluent cells to a mature epithelial layer is 7 and, thus, much larger than the substrate-dependent factor of 1.5.

### Role of Actin

A central theme of our work concerning cell mechanical and morphological properties is the role of reorganized actin filaments. Determining the underlying elements that influence cell mechanical properties is currently a highly active research area; though, actin filaments are emerging as a major factor. Our data, and literature reports^43,60^, suggest that cells possess a baseline stiffness of around 500 - 1000 Pa that may originate from the cytoplasm, the nucleus, the membrane, and cytoskeletal filaments, or a combination of these factors. Our data, and data in the literature, also suggest that enhanced stiffness – on top of the baseline stiffness – correlates with increased actin density. Fig. 4 shows that enhanced stiffness is measured in the region where cells abut, and the enhanced stiffness strongly co-localizes with dense actin filament bundles in confluent cells and in the epithelial layer cells. This agrees with the observations by Schierbaum et al.^55^. Enhanced stiffness is also seen in epithelial layer cells, in general, and it again strongly co-localizes with dense actin filament bundles. Several studies found that actin filaments play a major role in determining cell stiffness. Notably, treatment of cells with reagents, such as cytochalasin and latrunculin, which promote actin depolymerization, resulted in cells with significantly decreased stiffness.^29–33,58,60^ On the other hand, depolymerizing microtubules by treating cells with nocodazole had only marginal effects on cell stiffness and shape,^29,32–34^ suggesting that microtubules are only a minor contributor to cell stiffness. Furthermore, time-resolved measurements of cell stiffness on cells during monolayer formation showed that an increase in stiffness of the monolayer correlated with the assembly of adherens junctions but not desmosomes.^46^ This suggests intermediate filaments only play a minor role in cell stiffness. Additionally, diffuse cortical F-actin, as seen in the confocal images of single cell (Fig. 5a & d), does not appear to correlate with the enhanced stiffness, since the cortical F-actin does not vary noticeably with stiffness.

### Cell stiffness and cell geometric factors

The data in Fig. 3e & f regarding stiffness as a function of aspect ratio and as a function of cell area suggest that geometric factors correlate to cell stiffness for single cells but not for confluent cells. Lemmon et al. and Califano et al. demonstrated that the spreading area of a cell positively correlates with the magnitude of traction forces generated by the cell^63,64^. A balance between the traction force, cortical membrane tension and the cytoplasmic pressure is required to maintain cell shape. This implies that larger cells maintain a stronger tensional prestress. Since prestress strongly affects cell stiffness,^65–69^ it is expected that cells with a large spreading area are stiffer than cells with a small spreading area. This is consistent with our data showing that stiffness increases with cell area for single cells (Fig. 2f). Similar correlations were found in endothelial cells^9^. Cell area of single cells also correlated with their aspect ratio (Fig. S2), which is possibly due to a correlation between aspect ratio and cells stiffness (Fig. 2e). In contrast, in confluent cells, there was no correlation between those geometrical factors and stiffness. (Fig. 2f & g). This is probably because in confluent cells tensional prestress originates from actomyosin filaments linked to adherens junctions, and not from counterbalancing forces against traction forces, which depend on cell area. This view is supported by the observation that basal stress fibers and focal adhesions decrease as cells become confluent.

Even though we did not quantitatively measure the spreading area and height of the cells on the top of the epithelial layer, these cells appeared to be 3-4 times wider and thinner than the basal cells (Fig. 1, Fig. 5, Fig. S1). This extremely stretched and flattened morphology possibly indicates that much stronger tension was loaded over the cells on the top layer, making the cells recruit a large amount of actomyosin filaments at the adherens junctions to provide tension and maintain the cell shape, which, in turn, caused the dramatically enhanced stiffness.

## Conclusions

In summary, we found that HMECs possess significantly different mechanical properties along with different F-actin distributions as they transition from single to confluent to mature epithelial layer cells. The observed significant stiffening of cells in an epithelial layer is likely physiologically important, providing protection against external insults. There are likely additional intermediate transitioning states between confluent cells and mature epithelial cells, in which the stiffnesses progressively increase. Future work will focus on investigating the detailed mechanisms by which the cells on the top of the mature epithelial layer become flat and stiff. Our findings advance the understanding of breast ductal morphogenesis and mechanical homeostasis.

## Acknowledgements

This work was supported by an instrumentation grant for the AFM from the North Carolina Biotechnology Center (NCBC; 2014-IDG-1012), and grants from the Discover Institute and the NIH (1R15HL148842). We are grateful to Dr. Adam Hall, who is the PI of the NCBC instrumentation grant; to Heather Brown-Harding, who is the head of the Wake Downtown confocal microscope facility at Wake Forest University; to Pierre-Alexandre Vidi for helpful discussions; and to Amanda Smelser for help with cell culture work.

## Supplementary Information

**Figure S1.**
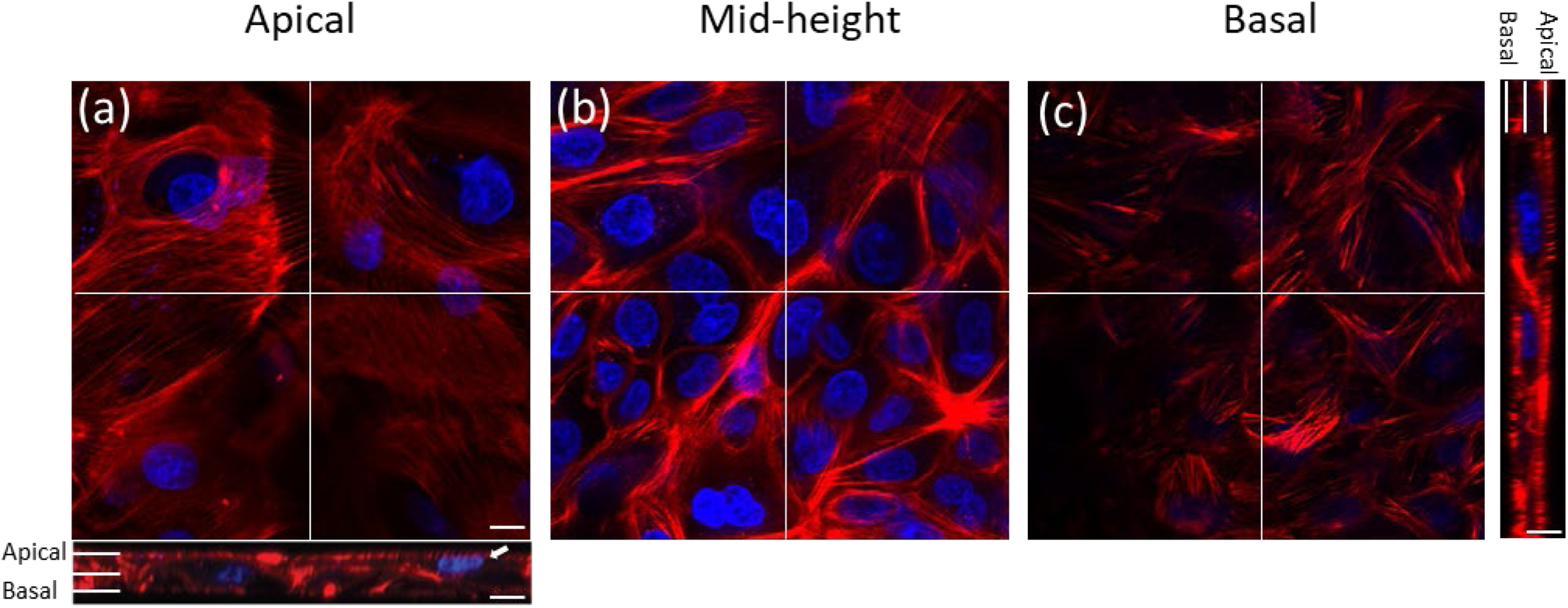
Confocal microscope images of F-actin (red) and nuclei (blue) of HMECs in an epithelial layer taken in (a) apical, (b) mid-height, and (c) basal focal planes. The white arrow in the horizontal cross-sectional view indicates a cell on the top layer of the epithelial layer. The scale bar is 10 µm.

**Figure S2.**
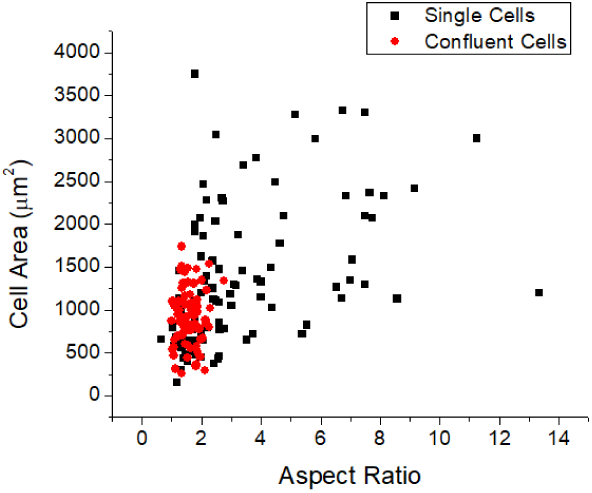
Cell area vs Aspect ratio plot for single cells and confluent cells.

